# Physiological and transcriptome analysis of indica rice ‘MH86’ overexpressing *OsSPL14*

**DOI:** 10.1101/496620

**Authors:** Ling Lian, Wei He, Qiuhua Cai, Hui Zhang, Chengrong Ren, Jun Ding, Huibin Xu, Yuelong Lin, Liyan Pan, Yidong Wei, Yongsheng Zhu, Yanmei Zheng, Linyan Wei, Hongguang Xie, Huaan Xie, Zonghua Wang, Jianfu Zhang

**Author notes:** Corresponding authors: Jianfu Zhang, Ph.D, Professor.

## Abstract

*OsSPL14*, identified as *IDEAL PLANT ARCHITECTURE1 (IPA1)* or *WEALTHY FARMER’S PANICLE (WFP)* gene, plays a critical role in regulating rice plant architecture. Here, the study showed that *OsSPL14*-overexpression transgenic rice plants had shorter growth period, short-narrow flag leaves, and thick-green leaves. Compared with wild type plant ‘MH86’, transgenic plants had higher chlorophyll a (Ca), chlorophyll b (Cb) and carotenoid (Cx) content at both seedling and maturity stage. Meanwhile, transcriptome analysis identified 473 up-regulated and 103 down-regulated genes in transgenic plant. The expression of differentially expressed genes (DEGs) involved in carotenoid biosynthesis, abscisic acid (ABA) metabolism and lignin biosynthesis increased significantly. Most of DEGs participated in “plant hormone signal transduction” and “starch and sucrose metabolism” are also up-regulated in transgenic plant. In addition, there were higher levels of ABA and gibberellin acid (GA_3_) in *OsSPL14*-overexpression transgenic plants. Moreover, the content of culm lignin, cellulose, silicon and potassium all increased dramatically. Thus, these results demonstrate that overexpression of *OsSPL14* has influence on leaf development, hormone level and culm composition in rice, which provide more insight into understanding the function of *OsSPL14*.

**Highlight:** *OsSPL14*-overexpression transgenic plants showed shorter growth period, short-narrow flag leaves, and thick-green leaves. Transcript profile, abscisic acid (ABA) and gibberellin acid (GA_3_) level, and culm composition also changed obviously.

## Introduction

*OsSPL14* is a member of the SQUAMOSA PROMOTER BINDING PROTEIN-LIKE (SPL) genes. SPL genes encode plant-specific transcription factors containing a highly conserved zinc ion–containing DNA binding domain, known as SQUAMOSA PROMOTER BINDING PROTEIN [SBP]-box (Yamasaki *et al*., 2004). SPB1 and SPB2 were the original SPL genes identified in *Antirrhinum majusas* that bind to the promoter of the floral meristem identity gene SQUAMOSA (Klein *et al*., 1996).

Many SPL genes contain a target site of miRNAs (microRNAs) including miR156/157, and the target sites are located in the coding regions or 3′ untranslated region (3′ UTR). In *Arabidopsis thaliana*, 10 of 16 SPL genes have been predicted to be targets of miR156 (Rhoades *et al*., 2002; Schwab R *et al*., 2005). *SPL3, SPL4*, and *SPL5* have a target site for the miR156 in their 3′ UTR and are strongly repressed by miR156. Overexpression of *SPL3* accelerated flowering, and plants expressing *SPL3* containing mutated miR156 target site or without *miR156* target site appeared earlier flowering and fewer leaves (Cardon *et al*., 1997; Wu and Poethig, 2006; Gandikota *et al*., 2007). Besides, paralogous genes *SPL9* and *SPL15* with miR156 target site involved in controlling the juvenile-to-adult phase transition, and *spl9 spl15* double mutants showed shortened plastochron, altered inflorescence architecture and enhanced branching(Schwarz *et al*., 2008; Wang *et al*., 2008). In addition, *SPL8* with no miRNA target sites affected megasporogenesis, trichome formation on sepals, and stamen filament elongation (Unte *et al*., 2003). Accordingly, a series of studies on *SPL* gene in *Arabidopsis thaliana* suggested that *AtSPL* genes mainly associated with plant development and flowering.

In rice, there are 19 SPL genes (*OsSPL*) that unevenly distributed in the genome. Among them, 11 *OsSPL* genes were putative targets of OsmiR156. *OsSPL* genes express in various tissues including root, leaf, stem, panicle, stamen, pistil, and most of them predominantly expressed in the young panicles (xie *et al*., 2006). Study on the liguleless mutant indicated that *OsLG1* encoding OsSPL8 participates in building the laminar joint between leaf blade and leaf sheath boundary, so as to control the development of ligule and auricle (Lee *et al*., 2007). *OsSPL14*, also known as the *IPA1/WFP* gene, is a homologous gene of the Arabidopsis SPL9/SPL15, and it is regulated by OsmiR156/OsmiR529 (Jeong *et al*., 2011; Yue *et al*., 2017). A point mutation in target site for OsmiR156 resulted in a significantly increased protein level of OsSPL14, which leads to reduced tiller number, increased grain number and grain weight, produced stout stems and improved lodging resistance (Jiao *et al*., 2010). Additionally, an epigenetic change in the *OsSPL14* promoter resulted in the same phenotype (Miura *et al*., 2010). Moreover, the three tandem repeats in the upstream regions of *IPA1/OsSPL14* enhanced an open chromatin configuration and reduced DNA methylation at *IPA1/OsSPL14* promoter, resulting in an optimal *OsSPL14* expression and ideal plant architecture with proper tiller number and large panicle (Zhang *et al*., 2017). While, the *ipa1* loss-of-function mutants generated by CRISPR/Cas9 had a dwarf phenotype with increased tiller number (Li *et al*., 2016). In addition, genome-wide expression profiling analysis suggested that IPA1/OsSPL14 could directly bind to the SBP-box target motif GTAC and regulated expression of TEOSINTE BRANCHED1 (TB1) and DENSE AND ERECTPANICLE1 (DEP1), so as to influence till number, plant height and panicle length (Lu *et al*., 2013). Then, according to *IPA1/OsSPL14* effect mentioned above, the studies on mechanism of gene expression regulation were carried out. The study indicated that a RING-finger E3 ligase named IPA1 INTERACTING PROTEIN1 (IPI1) could regulate the protein levels of *IPA1/OsSPL14* in different tissues. Interestingly, IPI1 promotes accumulation of IPA1/OsSPL14 in shoot apexes, but it promotes degradation of IPA1/OsSPL14 in panicles (Wang J *et al*., 2017). And the other research suggested that a human ovarian tumor domain-containing ubiquitin aldehyde-binding protein 1(OTUB1)-like deubiquitinase (*OsOTUB1*) could promote the degradation of IPA1/OsSPL14 indirectly (Wang S *et al*., 2017). Moreover, *DWARF 53* (*D53*), a key repressor of the strigolactone (SL) signaling pathway, suppressed the transcription of *IPA1/OsSPL14*. However, IPA1/OsSPL14 acted in the feedback regulation of *D53* expression (Song *et al*., 2017). The latest research indicated that phosphorylated IPA1/OsSPL14 activates expression of *WRKY45* and then enhanced disease resistance (Wang *et al*., 2018). In short, the previous studies revealed that IPA1/OsSPL14 played important role in controlling plant architecture and panicle architecture. Also, gene regulatory networks related to *IPA1/OsSPL14* are complex and the corresponding molecular mechanism requires further clarification.

In this study, we introduced *OsSPL14* into indica cultivar ‘MH86’, and obtained transgenic lines overexpressing *OsSPL14* with shorter growth period, short-narrow flag leaf, fewer tiller numbers, strong culm. Transcriptome analysis indicated that genes involved in carotenoid biosynthesis and lignin biosynthesis were up-regulated in transgenic plant. Levels of ABA and GA_3_ improved, and the content of culm lignin, cellulose, silicon and potassium increased obviously. Taken together, these discoveries supplement the function description of OsSPL14.

## Materials and Methods

### Generation of transgenic rice

The plasmid pCAMBIA1300-*OsSPL14* (Supplementary Fig.S1; *OsSPL14/IPA1:* GenBank GU136674.1, 7229 bp containing the promoter and coding region for mRNA) was introduced into mature rice (*Oryza sativa* L.indica cultivar ‘MH86’) embryos using Agrobacterium-mediated transformation as described by Datta (1997). The transformed cells were selected initially on NB medium (Sivamani *et al*., 1996) containing 50 mg/L hygromycin (SIGMA-ALORICH). After 3-4 selection cycles with a 14-day duration per cycle, surviving cell clusters were regenerated. The plants that were generated were selected further in the culture solution containing 20 mg/L hygromycin with a 16-h photoperiod at 26 °C for 4 weeks and then transferred to a greenhouse and grown with WT plants under natural sunlight.

Genomic DNA was prepared from the bialaphos-resistant plants by the Cetyltrimethylammonium bromide (CTAB) method and used as a template for Polymerase Chain Reaction (PCR) and southern blot analysis. The presence of the introduced gene in the regenerated plants was verified by PCR using the Hygromycin F/R primers. A total of 100 μg DNA per sample was digested overnight with *EcoRI* restriction enzyme at 37°C. The digested DNA was fractionated in 1% (w/v) TAE-agarose gels and then transferred to a Hybond-N nylon membrane (Amersham, Arlington Heights, IL, USA), according to the manufacturer’s instructions. A partial sequence of *Hygromycin* gene was isolated from the plasmid and labeled using an AlkPhos direct labeling kit (Amersham) in order to make the hybridization probes. Then the DNA fragments on the Hybond-N nylon membrane were hybridized with the *Hygromycin* probe.

Total RNA was isolated from PCR positive plants and WT using TRIzol reagent (Invitrogen, USA). First-strand cDNA was synthesized from 2 μg of total RNA using the RevertAidTM First Strand cDNA Synthesis Kit (Fermentas, LTU) following the manufacturer’s instructions. Reverse transcription polymerase chain reaction (RT-PCR) was performed using Hygromycin F/R and Actin150 F/R primers.

The transgenic plants were grown in transgenic experimental field. And stably inherited transgenic plants possessing two copies of the transgene were selected and used in this study.

### RNA sequencing and transcriptome analysis

Total RNA was isolated from one-month-old seedlings of transgenic plants and ‘MH86’ using TRIzol reagent. RNA purity and concentration were checked using NanoDrop microvolume spectrophotometer (Thermo ScientificNanoDrop Products, Waltham, MA, USA). And RNA integrity was quantified using Agilent 2100 Bioanalyzer® (Agilent Technologies, Palo Alto, Calif.). Then the enrichment of mRNA containing poly (A) was performed by Oligo (dT) magnetic beads. Subsequently, the mRNA was fragmentated and used to synthesize double-stranded cDNA (dsDNA). And the dsDNA was purified using AMPure XP beads (Beckmann Coul-ter, Pasadena, CA). Next, the “A” tail and paired-end adapters for sequencing were ligated to the end of dsDNA. Different sizes of fragments were selected using AMPure XP beads for PCR amplification to create the cDNA library. The concentration and insert size of the cDNA library were detected using Qubit2.0 and Agilent 2100, respectively. Meanwhile, the effective concentration of the cDNA library was accurately quantified using quantitative PCR. After complete testing, the cDNA library was sequenced using Illumina HiSeq2500 Sequencer. Low quality reads were filtered to get clean reads, then the clean reads were mapped to the reference genome (http://plants.ensembl.org/Oryza_indica/Info/Index) using TopHat2 and mapped reads were obtained.

Based on the location information of mapped reads, gene expression levels were calculated by cuffquant and cuffnorm components of cufflinks software using fragments per kilobase of transcript per million fragments mapped (FPKM) as measurement index. According to screening standard of fold Change (FC) ≥2 and false discovery rate (FDR) <0.01, differentially expressed genes (DEGs) analysis between *OsSPL14*-overexpression transgenic plants and wile type plant ‘MH86’ was perform by differential expression analysis for sequence count data (DEseq). The log2FC (fold changes) was used to indicate the significance of expression difference of each gene. The test corrections of P-values were performed using the Benjamini-Hochber False Discovery Rate method. The numbers of DEGs between samples were counted and plotted in Volcano plot.

For gene ontology (GO) enrichment analysis, DEGs were implemented in biological annotation system by Gene Ontology Consortium. And DEGs were classified according to biological process, molecular function and cellular component ontologies. The most enriched GO terms were conducted by KOBAS (2.0), and GO terms with FDR≤0.05 were considered as significantly enriched. The DEGs were subjected to search against Cluster of Orthologous Groups of proteins database to predict and classify their possible functions. Then the pathway enrichment and annotation analysis of DEGs were carried out by comparing to the KEGG (Kyoto Encyclopedia of Genes and Genomes) database. At last, taking pathway in KEGG database for unit, the DEGs significantly enriched metabolic pathways or signal transduction pathways were identified by hypergeometric test.

### Quantitative Real-time PCR (qRT-PCR)

To validate DEGs identified in the transcriptome analysis and analyze some genes from *OsSPL14* gene regulatory networks, qRT-PCR was conducted on selected genes. For qRT-PCR, cDNA was generated from the previously collected RNA using the RevertAidTM First Strand cDNA Synthesis Kit (Fermentas, LTU). The qRT-PCR was performed on a 7500 real-time PCR system (Applied Biosystems) with the Fast Start Universal SYBR Green Master system (Roche, USA), following the manufacturer’s protocol. Relative quantification of gene expression was performed using actin gene expression as a reference. The relative quantitative method (ΔΔCT) was used to evaluate the quantitative variation in the examined replicates. Primers used in qRT-PCR are listed in Supplementary Table S1. Additionally, TB F/R and DEP F/R primers, D53 F/R primers, and D17 F/R, D14 F/R, D3 F/R primers were referenced to Lu *et al*. (2013), Song *et al*. (2017), Sun *et al*. (2014), respectively.

### Assay of abscisic acid (ABA), jasmonate acid (JA) and gibberellin acid (GA_3_)

One-month-old seedlings and plants at tillering stage were collected to reserve. And ABA, JA, GA contents were detected by high performance liquid chromatography (HPLC, Agilent 1100).

For extraction of ABA, 0.15 g plant material was manually ground in liquid nitrogen. Samples were taken in 1 mL ABA extraction reagent (Suzhou Comin Biotechnology Co. Ltd) at 4 °C for overnight. Samples were centrifuged for 10 min at 8000 ×g, and the residue was extracted with 0.5 mL ABA extraction reagent for 2 hours again. After 2 times centrifugation, the supernatant was collected to be dried with 40 °C nitrogen until containing no organic phase. Then 0.5 mL petroleum ether was added to extract and decolorize at 60-90 °C for three times. The samples in the lower layer were collected to be dried with 40 °C nitrogen, then concussed and dissolved in 0.5 mL mobile phase containing acetic acid and methanol. Next, the samples were filtered through minisart filters for HPLC analysis. Chromatographic experiments were performed on Kromasil C18 column (250mm*4.6mm, 5 μm) with a mixture acetic acid-methanol as the mobile phase, a flow rate of 0.8 mL/min, a detection wavelength at 254 nm, and column temperature at 30 °C.

JA was extracted using 2 g plant material in Eppendorf tube containing JA extraction reagent (Suzhou Comin Biotechnology Co. Ltd). After 30 min sonication, samples were centrifuged for 10 min at 8000 ×g. The supernatant was collected to be dried with nitrogen in ice-water bath. Then the samples were dissolved in 0.5 mL mobile phase of methanol: 0.1% formic acid (65: 35) and filtered for HPLC analysis. Chromatographic experiments were performed with a flow rate of 0.8 mL/min, a detection wavelength at 210 nm, and column temperature at 35 °C.

GA was extracted using 1.5 g plant material in 1 mL GA extraction reagent (Suzhou Comin Biotechnology Co. Ltd) at 4 °C for overnight. After centrifugation, the supernatant was collected to be dried. And 0.5 mL petroleum ether was added to extract and decolorize at 60-90 °C for three times. Next, the samples were extracted in reagent and dried with nitrogen. Then the samples were dissolved in 0.5 mL mobile phase of 0.1% acetic acid: methanol (60: 40) and filtered for HPLC analysis. Chromatographic experiments were performed Kromasil C18 column with a flow rate of 0.8 mL/min, a detection wavelength at 254 nm, and column temperature at 30 °C.

### Scanning electron microscopy (SEM) analysis

At heading stage, a 0.3-cm section was cut with a scalpel in the middle of the third internode and placed in 3% glutaraldehyde for fixing 8-10 h. After rinsed with phosphate buffer (0.1mol/L, PH7.2), samples were fixed with 1% osmic acid for 1h. Samples were rinsed with phosphate buffer, then they were dehydrated in a grade of ethanol series (35%, 50%, 60%, 70%, 95%, 100%, 100%) for 1h per grade. Next, samples were transferred into isoamyl acetate and dried using CO_2_ as transitional fluid. Finally, the samples were mounted onto aluminum stubs and coated with carbon gold, then observed with 3500N scanning electron microscope (Hitachi, Tokyo, Japan) to collect images.

### Composition analysis of culm

At the mature stage, plants culm was collected and placed in an oven at 105°C for 30 min, then transferred to 65°C until a constant weight was achieved. Then the plant samples were crushed in a grinder to make powder for test.

For determination of cellulose and lignin content, 1 g powder sample was treated with 100 mL 3% sodium dodecyl sulfate containing three drop of decalin and 0.5 g anhydrous sodium sulfite in beaker. And the sample solution was heated to boiling for 1 h. After filtration, the precipitate was washed with hot water and acetone, then dried 3 h at 105 °C to weigh. Next, the precipitate was heated and digested with 100 mL 2% cetyltrimethylammonium bromide. The sample solution was filtered and the second precipitate was obtained, then washed and dried to weigh. Finally, the second precipitate was treated with 72% sulfate for 3 h, then desiccated to weigh. The numerical calculation of cellulose and lignin content was reference to method described in GB/T 13885-2003/ISO 6869:2000; 2000.

For measurement of potassium content, 2 g powder sample was digested with 30mL nitric acid and perchloric acid (5:1) overnight. Then the sample solution was treated by heating digestion method to be clear. After forging, deionized water was added to a constant volume of 100 mL and filtered for detection. According to the standard curves of potassium, potassium content was measured with atomic absorption spectrometry (AAS) at 766.5 nm (GB/T 13885-2003/ISO 6869:2000; 2000).

1 g powder sample sifted by 60-mesh sieve was using to measure silicon (Si) content. 15 mL concentrated sulfuric acid was gradually added to the powder sample in beaker. 40 min later, 50 mL concentrated nitric acid was gradually added to it and slow heating to precipitate. Then sodium hydroxide was added to dissolve the precipitate, and the silicon (Si) content of culm was determined using the silicon molybdenum blue colorimetric method (Tong *et al*., 2005; Dai *et al*., 2005).

## Results

### Phenotype of MH86 over-expressing *OsSPL14*

We introduced “pCAMBIA1300-*OsSPL14*” into indica cultivar MH86 using agrobacterium-mediated transformation and generated the *OsSPL14*-overexpression transgenic plants (OE: *OsSPL14;* Supplementary Fig. S2). Remarkably, the growth period of OE: *OsSPL14* transgenic plants shortened by 5-15 days depended on the growing areas (Fig. 1A). OE: *OsSPL14* transgenic plants had shorter and narrower flag leaves. Compared to that of wild type plant ‘MH86’ (WT), the flag leaf length and width of OE: *OsSPL14* transgenic plant decreased by about one third and one quarter respectively (Fig. 1B, C, D). Well consistent with previous report (Jiao *et al*., 2010), OE: *OsSPL14* transgenic plants showed decreased tiller number and strong culm (Fig. 1E). The culm diameter and culm wall thickness of transgenic plant increased by about 24% and 20% respectively (Fig. 1F, G). However, the height of OE: *OsSPL14* transgenic plants was not significantly different from that of ‘MH86’ (Fig. 1H). In addition, it is obvious that OE: *OsSPL14* transgenic plants appeared greener, especially at the maturity stage. Also, the leaves of transgenic plants were thicker than that of ‘MH86’. And the result of measurement indicated that the chlorophyll a (Ca), chlorophyll b (Cb) and carotenoid (Cx) content of leaves increased obviously at seedling stage and maturity stage (Fig. 1I, J).

**Fig.1.**
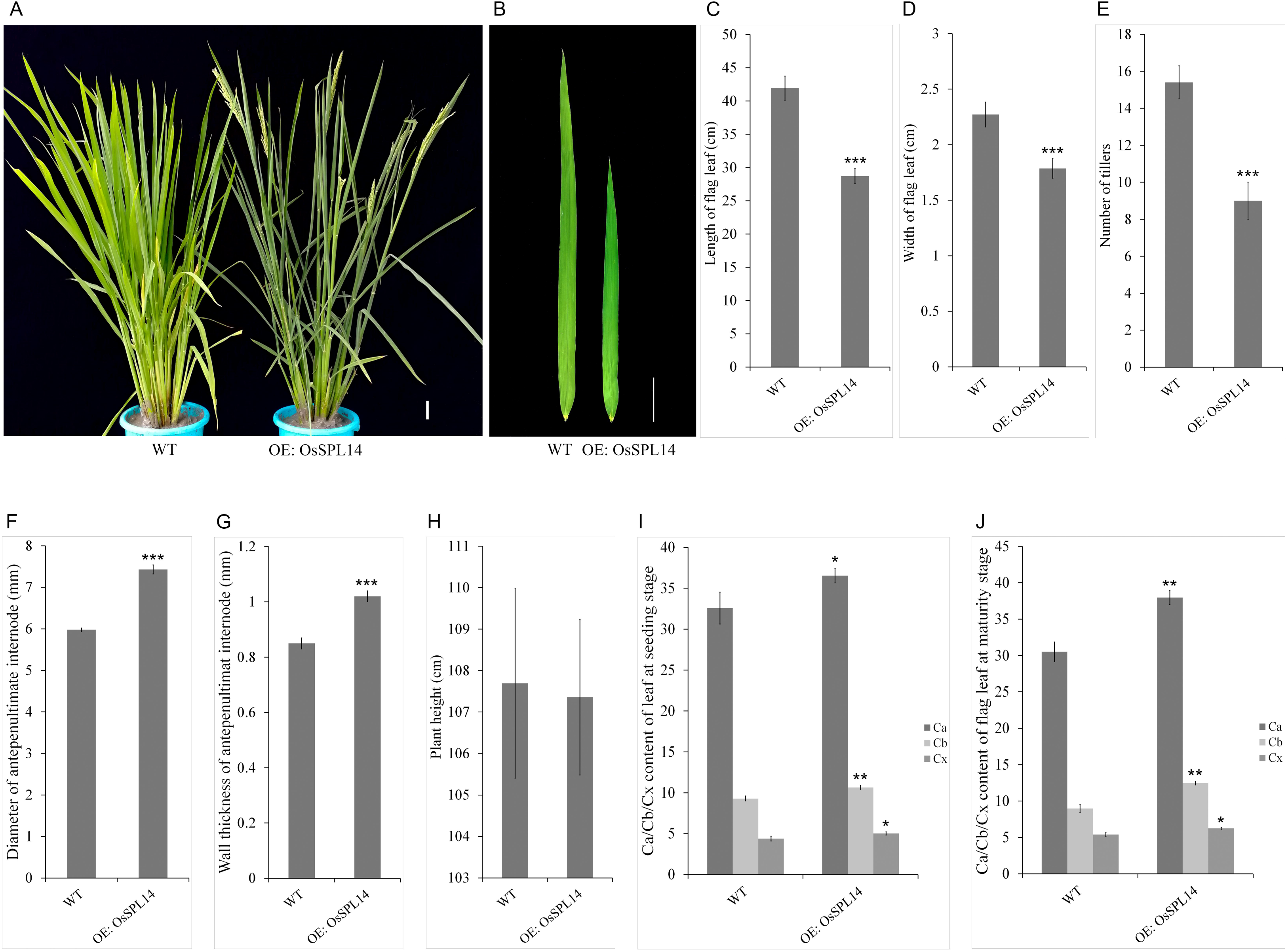
Analysis of plant phenotype. (A) The plant architecture of wild plant MH86 and OE: *OsSPL14* transgenic plant. (B) Leaf morphological character. (C) Length of flag leaf. (D) Width of flag leaf. (E) Number of tillers. (F) Diameter of antepenultimate internode. (G) Wall thickness of antepenultimat internode. (H) Plant height. (I) Chlorophyll a, Chlorophyll b and carotenoid content of leaf at seeding stage. (J) Chlorophyll a, Chlorophyll b and carotenoid content of flag leaf at maturity stage. Bar=5 cm. (*P≤0.05, **P≤0.01, ***P≤0.001).

### Comparative analysis of transcriptome profiling

To investigate the impact of *OsSPL14* overexpression on the rice transcriptome, we conducted an RNA sequencing experiment using OE: *OsSPL14* transgenic line and ‘MH86’ (Control). Differentially expressed genes (DEGs) was identified by screening standard thatfold Change (FC) ≥2 and false discovery rate (FDR) <0.01. And the analysis revealed a total of 576 differentially expressed genes containing 473 up-regulated genes and 103 down-regulated genes (Fig. 2A).

**Fig.2.**
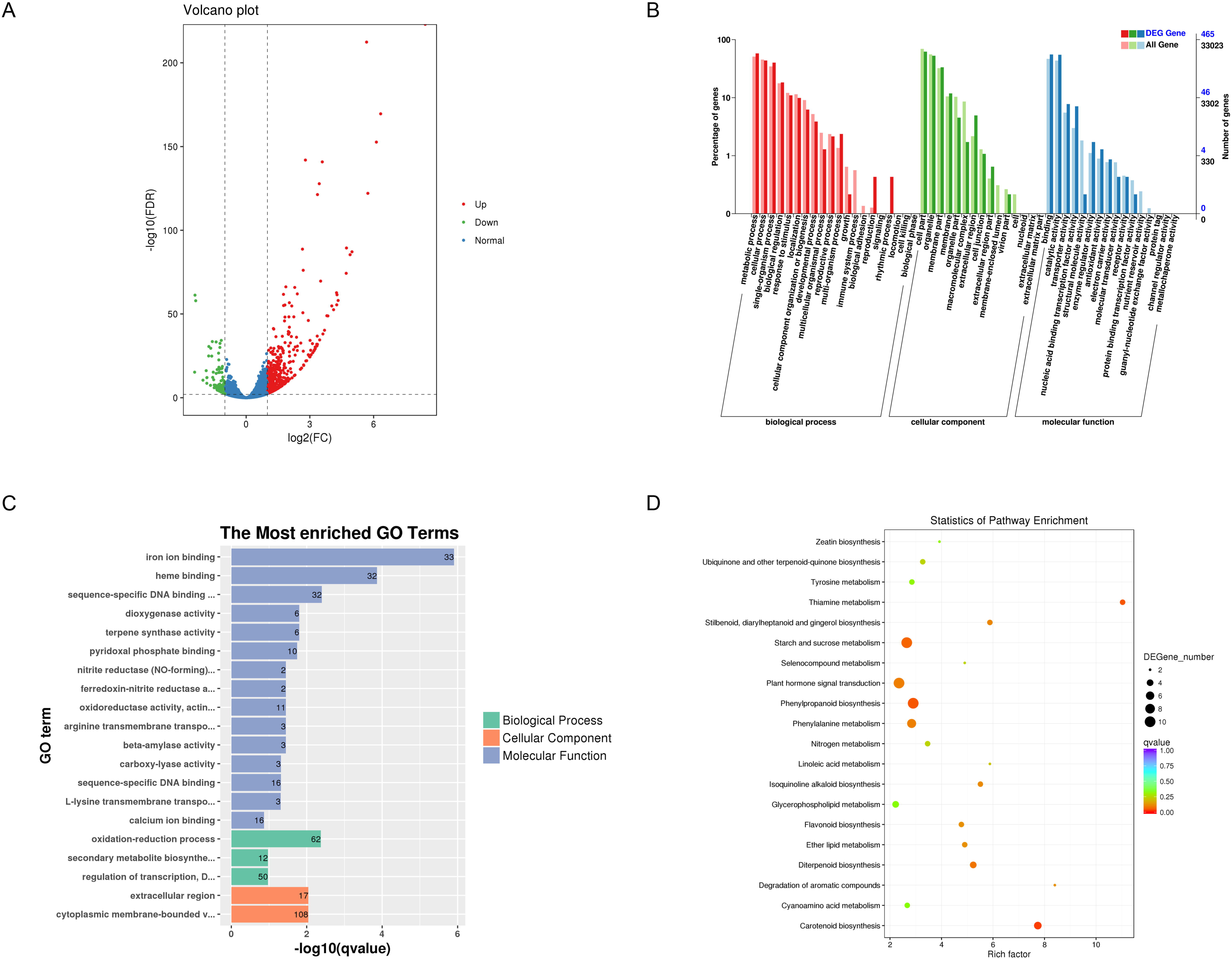
Transcriptome analysis. (A) Volcano plot of DEGs. Larger absolute value of abscissa means greater different multiple, and larger ordinate value means more significantly differential expression. Each dot represents a gene, in detail, green dots represent down-regulated genes, red dots represent up-regulated genes, and blue dots represent non-differentiated genes. (B) Gene ontology classifications of DEGs. The abscissa is GO classification, the left side of the ordinate is the percentage of genes, and the right side is the number of genes. (C) The most enriched GO terms. The ordinate is enriched GO term, and the abscissa is the q-value. The digitals on the column represent the number of differentially expressed genes. Different colors represent biological processes, cellular components and molecular functions respectively. (D) Scatter diagram of KEGG enrichment. Longitudinal axis represents pathway name, and lateral axis represents rich factor. The size of the point indicates the number of differentially expressed genes in the pathway, and the color of the point corresponds to a different Qvalue range.

Subsequently, gene ontology (GO) was used to classify the functions of the DEGs. The DEGs were categorized into three major GO categories including biological process, cellular component and molecular function, which totally contained 51 function groups (Fig. 2B). Meanwhile, the most enriched GO terms were conducted by KOBAS (2.0) with FDR≤0.05, and it showed that DEGs were grouped into 20 different GO terms, including 15 terms for molecular function, 3 terms for biological process and 2 terms for cellular component (Fig. 2C). It implied that DEGs annotated with GO terms were mainly related to “ion binding”, “heme binding”, “sequence-specific DNA binding”, “oxidation-reduction process”, “regulation of transcription” and “cytoplasmic membrane-bounded”.

Moreover in order to further understand the biological functions of the DEGs, we mapped the DEGs in kyoto encyclopedia of genes and genomes (KEGG) database and obtained 20 KEGG pathways that the most remarkably enriched (Fig. 2D; Supplement Table S2). And according to enrichment factor, Q value and the number of enriched genes in corresponding pathway, five KEGG pathways were selected to be annotated in details, which showed that most of the DEGs in these pathways were up-regulated (Table 1). In particular, the five DEGs in “Carotenoid biosynthesis” were all up-regulated. Among them, “BGIOSGA011548” encoding beta-carotene hydroxylase 2 (DSM2) and “BGIOSGA029294” encoding Phytoene synthase (PSY) are the important genes involved in carotenoid biosynthesis. Because carotenoid is the precursor of abscisic acid (ABA), the other genes “BGIOSGA025169” and “BGIOSGA013214” encoding 9-cis-epoxycarotenoid dioxygenase (NCED), “BGIOSGA029635” encoding Abscisic acid 8′-hydroxylase 3 (8′-OH-ABA3) also participate in ABA metabolism. The up-regulated DEGs in “Phenylpropanoid biosynthesis” were “BGIOSGA005999” encoding phenylalanine ammonia-lyase (PAL), “BGIOSGA006502” and “BGIOSGA008177” encoding trans-cinnamate 4-monooxygenase/cinnamate-4-hydroxylase (C4H), “BGIOSGA026897” encoding “4-coumarate-CoA ligase” (4CL), which also are key enzyme genes for lignin just because lignin is a phenylpropanoid monomer polymer (Zhong and Ye, 2015). The pathway showed the highest enrichment factor was “Thiamine metabolism”, in which the up-regulated DEG was “BGIOSGA025304” encoding 1-deoxy-D-xylulose-5-phosphate synthase 2 (DXS). DXS is the important enzyme involved in terpenoid anabolism, such as the biosynthesis of chlorophylls, tocopherols (VE), carotenoids, ABA, gibberellin (GAs), and so on (Aharoni *et al*., 2005; Estévez *et al*., 2001). Additionally, the DEGs in “Plant hormone signal transduction” are related to jasmonate acid (JA), ethylene and ABA signal transduction. The DEGs in “Starch and sucrose metabolism” are mainly related to sucrose, starch, trehalose, fructan and pectin metabolism.

**Table 1.** Annotation of five KEGG-enrichment pathways.

Gene ID in *EnsemblPlants* \*2: The Rice Annotation Project Database
| KEGG pathway | Gene ID<sup>*1</sup> | Gene ID in RAP<sup>*2</sup> | Regulation | Log2FC | Anno |
| --- | --- | --- | --- | --- | --- |
| **Carotenoid biosynthesis** | BGIOSGA011548 | Os03g0125100 | Up | 1.28 | Beta-ε |
|  | BGIOSGA029294 | Os09g0555500 | Up | 1.40 | Phyto |
| | BGIOSGA013214 | Os03g0645900 | Up | 1.57 | 9-cis-ε |
| | BGIOSGA025169 | Os07g0154100 | Up | 1.51 | 9-cis-ε |
|  | BGIOSGA029635 | Os09g0457100 | Up | 1.10 | Absci |
| **Phenylpropanoid biosynthesis** | BGIOSGA005999 | Os02g0626532 | Up | 1.01 | Pheny |
|  | BGIOSGA006502 | Os02g0466900 | Up | 1.08 | Trans- |
|  | BGIOSGA008177 | Os02g0467600 | Up | 1.41 | Trans- |
|  | BGIOSGA026897 | Os08g0448000 | Up | 1.10 | 4-cou |
|  | BGIOSGA014207 | Os04g0651000 | Down | -1.13 | class l |
|  | BGIOSGA022766 | Os06g0306300 | Down | -1.62 | class l |
|  | BGIOSGA027874 | Os08g0113000 | Down | -1.21 | class l |
| **Thiamine metabolism** | BGIOSGA025304 | Os07g0190000 | Up | 1.13 | 1-deo |
|  | BGIOSGA013327 | Os03g0679700 | Down | -1.23 | Phosp |
|  | BGIOSGA025849 | Os07g0529600 | Down | -1.17 | Cyste |
| **Plant hormone signal transduction** | BGIOSGA003839 | Os01g0583100 | Up | 1.24 | Protei |
|  | BGIOSGA011032 | Os03g0268600 | Up | 1.05 | Protei |
|  | BGIOSGA015611 | Os04g0167875 | Up | 1.54 | Protei |
|  | BGIOSGA017925 | Os05g0457300 | Up | 1.06 | Protei |
|  | BGIOSGA030517 | Os09g0325700 | Up | 1.61 | Protei |
|  | BGIOSGA009392 | Os03g0860100 | Up | 1.04 | Ethyle |
|  | BGIOSGA016308 | Os04g0395800 | Up | 2.41 | Jasmc |
|  | BGIOSGA011983 | Os03g0180900 | Up | 1.12 | Jasmc |
|  | BGIOSGA029689 | Os09g0439200 | Up | 1.80 | Jasmc |
|  | BGIOSGA010919 | Os03g0297600 | Down | -1.36 | Absci |
| **Starch and sucrose metabolism** | BGIOSGA007345 | Os02g0106100 | Up | 2.03 | Beta-δ |
| | BGIOSGA011829 | Os03g0141200 | Up | 1.41 | Beta-α |
|  | BGIOSGA031437 | Os10g0553300 | Up | 1.02 | Treha |
|  | BGIOSGA031535 | Os10g0521000 | Up | 1.16 | alpha, |
|  | BGIOSGA000847 | Os01g0743200 | Up | 1.15 | Pectin |
|  | BGIOSGA028767 | Os08g0450100 | Up | 2.01 | Pectin |
| | BGIOSGA007188 | Os02g0131400 | Up | 1.71 | Beta-δ |
| | BGIOSGA014898 | Os04g0474300 | Up | 1.14 | Beta-δ |
|  | BGIOSGA019700 | Os05g0361500 | Down | -1.14 | Pectin |
| | BGIOSGA032689 | Os10g0323500 | Down | -1.07 | Beta-δ |

### Gene expression analysis by qRT-PCR

Firstly, we analyzed the expression of *OsSPL14* in OE: *OsSPL14* transgenic lines. As expected, the expression of *OsSPL14* increased obviously in the detected transgenic lines (Fig. 3A). To confirm DEGs identified in the transcriptome analysis, several genes from the five KEGG pathways selected in last part were analyzed by qRT-PCR. And the result suggested that the expression of 15 up-regulated DEGs were higher in OE: *OsSPL14* transgenic plants than that in ‘MH86’ (Fig. 3B). Meanwhile, the expression of five down-regulated DEGs were lower in OE: *OsSPL14* transgenic plants than that in ‘MH86’ (Fig. 3C). In short, the expression profiles of DEGs analyzed by qRT-PCR were well consistent with transcriptome analysis.

**Fig.3.**
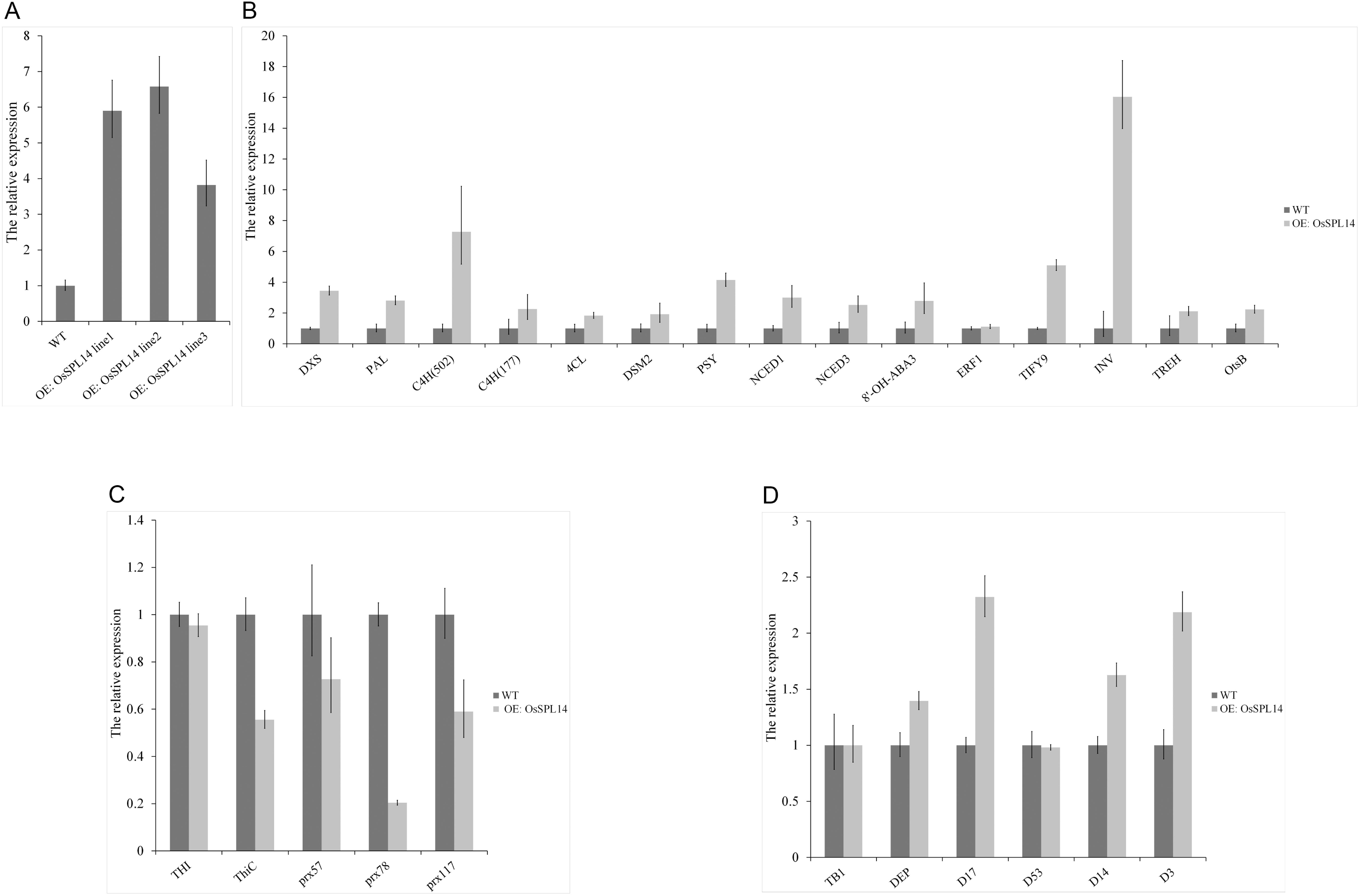
qRT-PCR analysis. (A) Expression analysis *of OsSPL14* in WT and OE: OsSPL14 transgenic lines. (B) Expression patterns of 15 up-regulated genes were validated by qRT-PCR. C4H (502): BGIOSGA006502; C4H (177): BGIOSGA008177. (C) Expression patterns of 5down-regulated genes were validated by qRT-PCR. (D) The expression profiles of *TB1, DEP1, D17, D53, D14* and *D3*.

According to previous research, OsSPL14 regulated expression of TB1 and DEP1, and OsSPL14 may act with D53 to mediate the Strigolactone (SL)-regulated tiller development (Lu *et al*., 2013; Song *et al*., 2017). So the expression profiles of *TB1*, *DEP1* and some genes related to SL biosynthesis or SL signal transduction were also analyzed by qRT-PCR. The result indicated that *TB1* had no obvious change, while *DEP1* was up-regulated slightly in OE: *OsSPL14* transgenic plants. In addition, compared with ‘MH86’, *DWARF* 17 (*D17*) involved in SL biosynthesis was up-regulated in OE: *OsSPL14* transgenic plants. *DWARF* 14 (*D14*) and *DWARF* 3 (*D3*) involved in SL signal transduction were up-regulated in transgenic plants (Fig. 3D). These results suggested that overexpression of OsSPL14 could indeed influenced on the expression of several genes involved in plant architecture.

### Changes in ABA, JA and GA_3_ content

In contrast with that of ‘MH86’, the phenotype of OE: *OsSPL14* transgenic plant changed greatly. Coincidentally, according to transcriptome analysis, it is remarkable that the expression of several genes in carotenoid biosynthesis, ABA metabolism, terpenoid anabolism and plant hormone signal transduction pathways are up-regulated. Thus, we are particularly intrigued that whether the content of relevant plant hormones changed in OE: *OsSPL14* transgenic plants. To ascertain it, we randomly selected two OE: *OsSPL14* transgenic lines to measure ABA, JA and GA3 content of OE: *OsSPL14* transgenic plants at seeding stage and tillering stage. Compared to that of ‘MH86’, ABA content of OE: *OsSPL14* transgenic plants increased obviously at seeding stage, by 18% and 23% in two transgenic lines respectively. The ABA content of transgenic plant increased slightly at tillering stage (Fig. 4A). However, JA content has no change in transgenic plant at seeding stage and decreased slightly in one transgenic line at the tillering stage (Fig. 4B). In contrast, the GA3 content of transgenic plants increased dramatically at the seeding stage, which was about three times that of ‘MH86’. Though both the GA3 content of transgenic lines and ‘MH86’ decreased significantly at tillering stage, the GA3 content of transgenic lines still increased by 56% and 27% compared to that of ‘MH86’ (Fig. 4C). Together, the observations indicated that there are higher level of ABA and GA_3_ in OE: *OsSPL14* transgenic plants than that in ‘MH86’.

**Fig.4.**
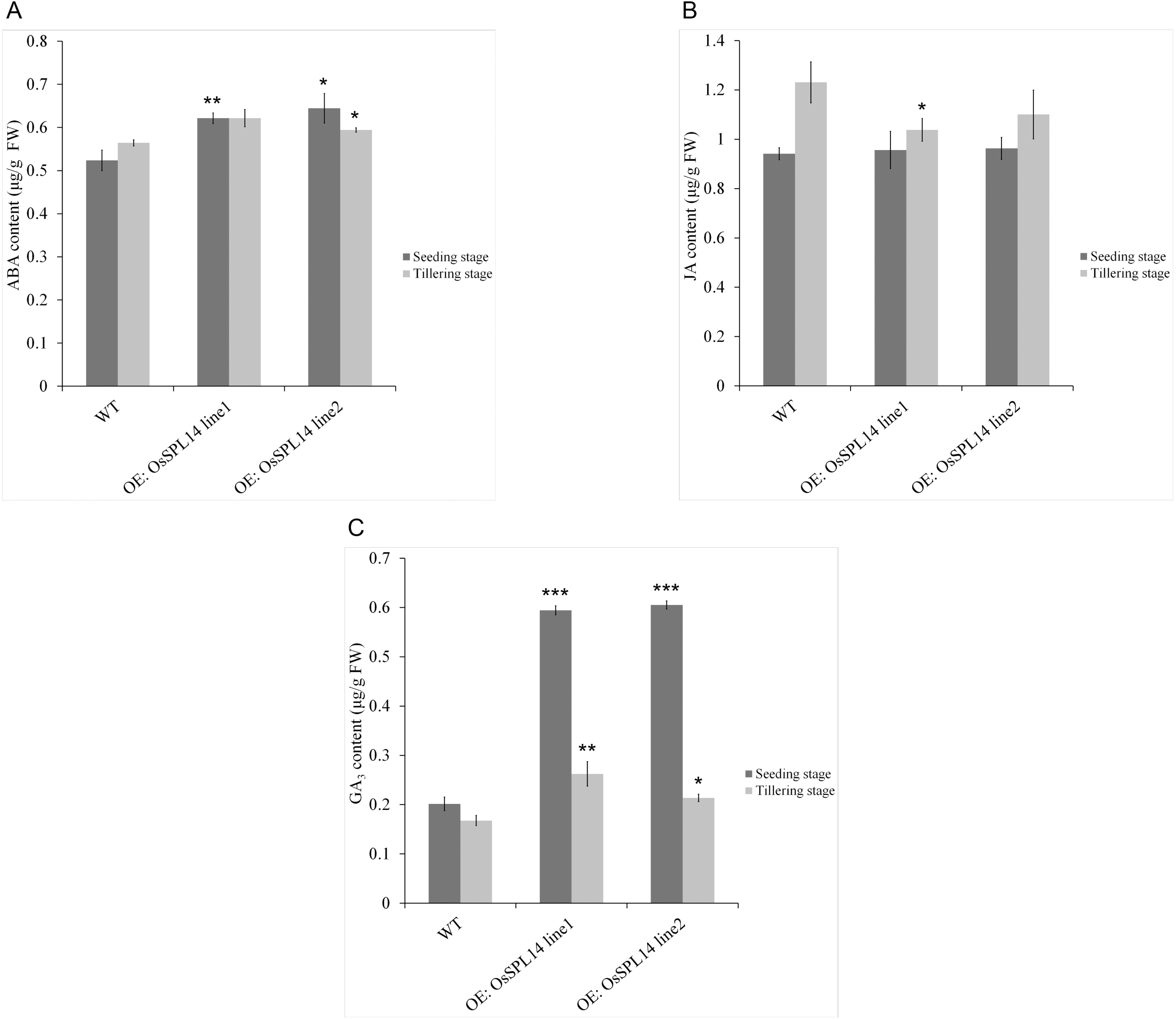
ABA, JA and GA_3_ content analysis. (A) ABA content at seeding stage and tillering stage. (B) JA content at seeding stage and tillering stage. (C) GA_3_ content at seeding stage and tillering stage. (*P≤0.05, **P ≤0.01, *** P≤0.001).

### Scanning and composition analysis of culm

One obvious characteristic of OE: OsSPL14 transgenic plants is their strong culm. Consequently, the internal organizational structure of culm was observed by scanning electron microscopy (SEM). The observation showed that OE: OsSPL14 transgenic plants had more small vascular bundles and sclerenchyma cells, and also had larger big vascular bundles. Moreover, it was obvious that the lignified degree of cortical cell was greater in OE: OsSPL14 transgenic plant (Fig. 5A).

**Fig.5.**
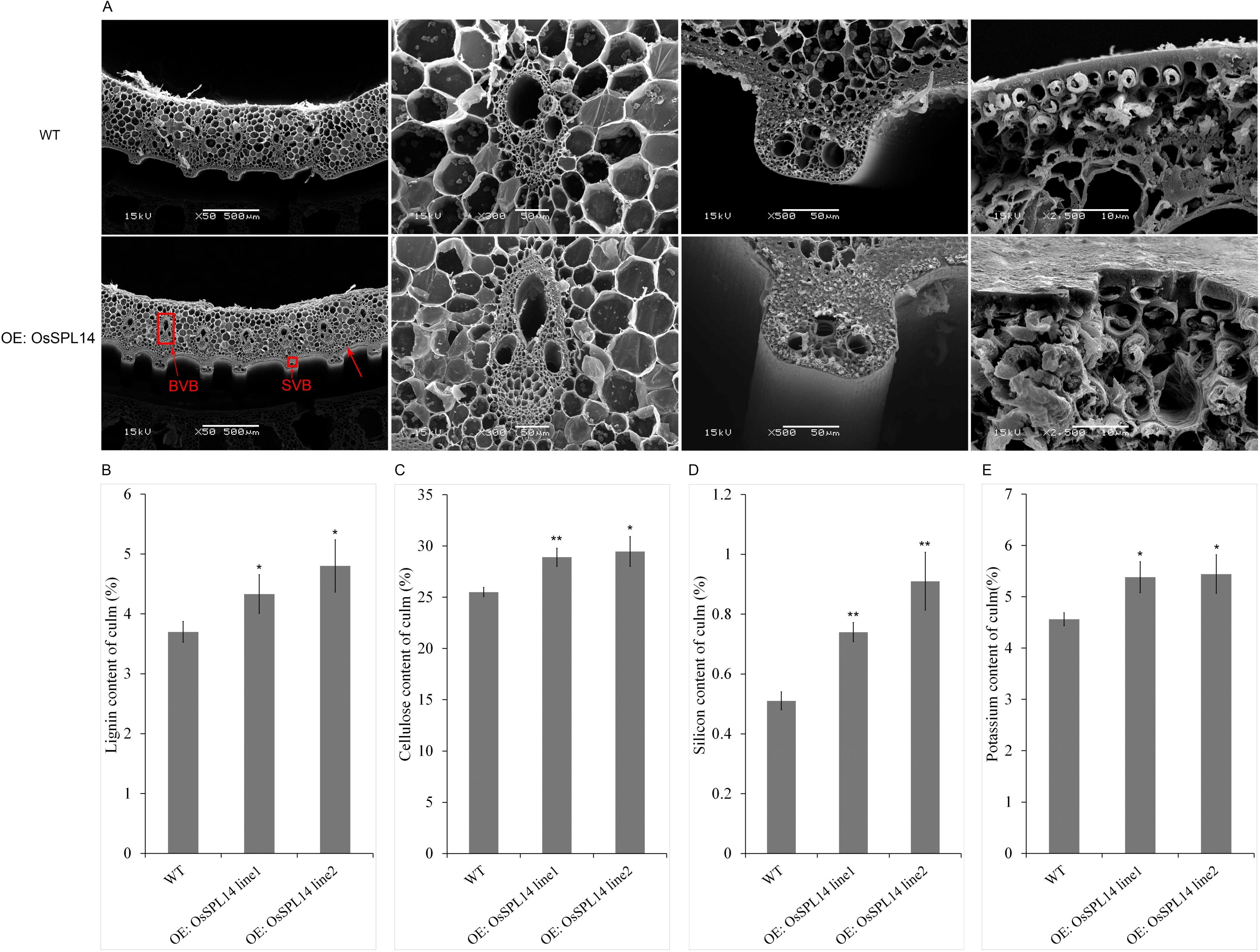
Scanning and chemical components analysis of culm. (A) Electron scanning microscopy analysis. BVB: Big vascular bundle; SVB: Small vascular bundles; SC: sclerenchyma cell. (B) Lignin content of culm. (C) Cellulose content of culm. (D) Silicon content of culm. (E) Potassium content of culm.

Based on transcriptome analysis in this article and the changes in internal organizational structure of culm, it is noteworthy that the expression profiles of the key genes involved in lignin biosynthesis and carbohydrates metabolism changed significantly. Subsequently, we hypothesized that culm composition of OE: OsSPL14 transgenic plant was likely to change. To test this prediction, we measured the lignin, cellulose, silicon and potassium content of culm. In contrast with that of ‘MH86’, the lignin content of OE: OsSPL14 transgenic plant increased distinctly, by 17% and 30% in the two lines respectively (Fig. 5B). And the cellulose content of transgenic lines also increased by 13% and 16% (Fig. 5C). Remarkably, the silicon content of transgenic plant increased dramatically by 45% and 78% compared to that of ‘MH86’ (Fig. 5D). In addition, the potassium content also increased obviously by 18% and 19% respectively (Fig. 5E). Evidently, overexpression of OsSPL14 in rice would cause great changes in culm character and composition. Besides, it is well known that culm composition is closely related to culm mechanical strength, which may likely explain the previously findings that the culm mechanical strength of the NIL OsSPL14^ipa1^ plant with high expression of OsSPL14 was significantly increased (Jiao *et al*., 2010).

### Grain quality analysis

It is clear that transgenic lines overexpressing OsSPL14 have large panicle, more branching and more grains per panicle (Jiao *et al*., 2010; Miura *et al*., 2010). Here, we performed grain quality analysis of OE: OsSPL14 transgenic plant. The result indicated that brown rice rate, milled rice rate and gel consistency had no obvious change in transgenic plant (Fig. 6A, B, C). While, head rice rate and grain length of OE: *OsSPL14* transgenic plant decreased slightly (Fig6. D, E), and the gelatinization temperature elevated slightly (Fig6. F). The amylose content of transgenic plant decreased obviously (Fig6. G). However, the chalkiness ratio of OE: OsSPL14 transgenic plant was one fold higher than that of ‘MH86’ (Fig6. H). Meanwhile, the chalkiness degree was 2.2 times that of ‘MH86’ (Fig6. I). The higher chalkiness ratio and chalkiness degree induced the decreased transparency (Fig6. J). Therefore, it seemed that some grain quality characters of OE: OsSPL14 transgenic plant changed to a certain extent.

**Fig.6.**
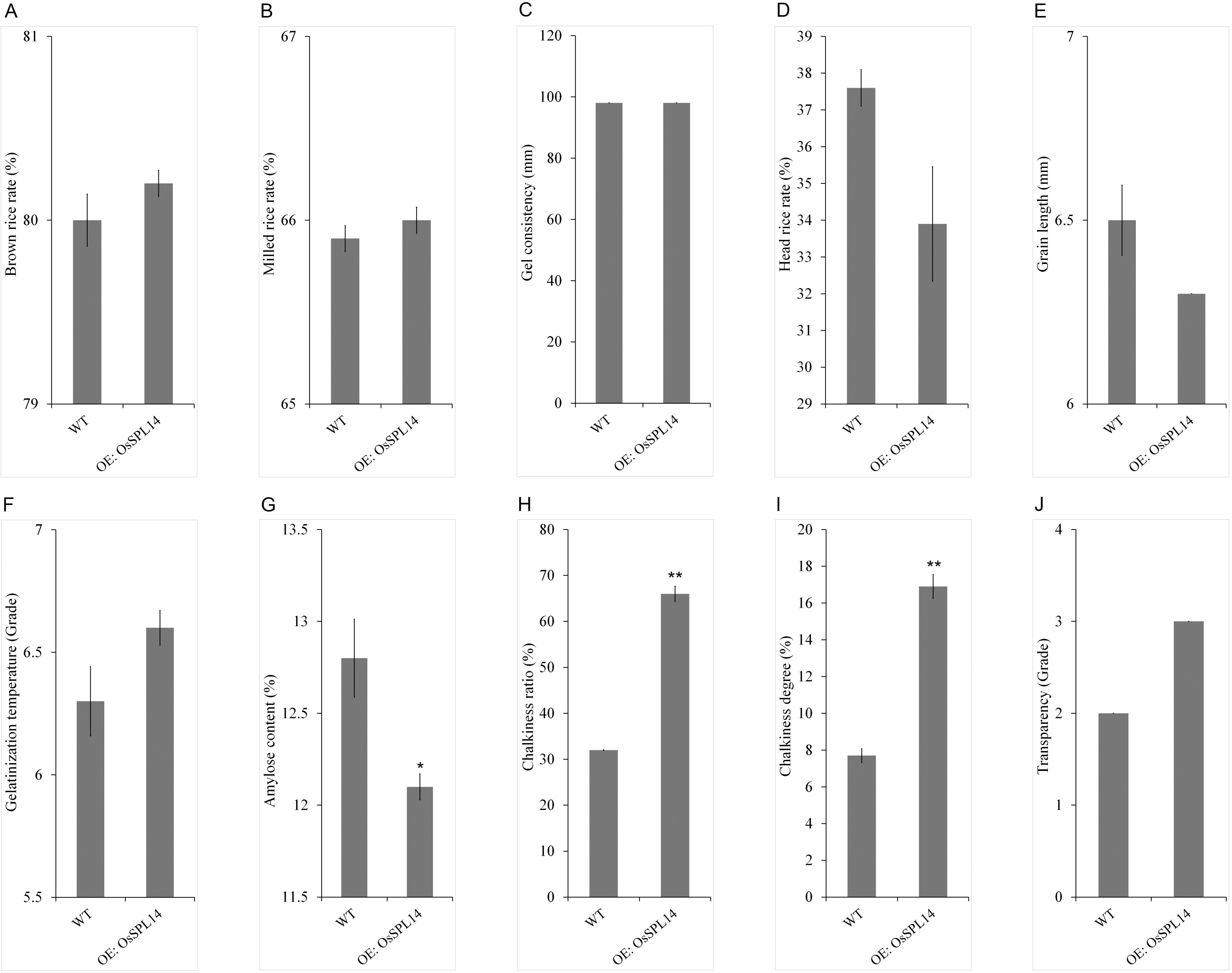
Grain quality analysis. (A) Brown rice rate. (B) Milled rice rate. (C) Gel consistency. (D) Head rice rate. (E) Grain length. (F) Gelatinization temperature. (G) Amylose content. (H) Chalkiness ratio. (I) Chalkiness degree. (J) Transparency. (*P≤0.05, **P≤0.01)

## Discussion

Os SPL14, as a transcription factor, plays an important role in rice plant architecture. The typical characteristics of transgenic plant overexpressing *OsSPL14* were reduced tiller number and strong culm. Expect for these characteristics, we observed that OE: *OsSPL14* transgenic plant have shorter growth period. It is consistent with previous research that overexpression of SPL3 caused early flowering and SPL9/SPL15 controlled the juvenile-to-adult phase transition in *Arabidopsis thaliana* (Cardon *et al*., 1997; Schwarz *et al*., 2008). Further study suggested that SPL3 impacted on flowering through directly activating plant-specific transcription factor LEAFY (LFY), MADS domain protein FRUITFULL (FUL) and APETALA1 (AP1), which controlled meristem identity (MI) transition (Yamaguchi *et al*., 2009). A previous study showed that the SPL genes affect flowering as downstream targets of CONSTANS (CO) and flowering time regulator (FT) at the shoot apex (Schmid *et al*., 2003). But the later study indicated that SPLs control flowering in a separate endogenous flowering pathway differed from familiar FT/FD (FD: bZIP transcription factor) flowering model, and the miR156/SPL module promoted flowering in the absence of photoperiodic cues (Wang *et al*., 2009). Thus, the molecular mechanisms of SPLs pathway controlling flowering are not fully understood, and there is hardly research on the molecular mechanisms in rice.

It is obvious that OE: *OsSPL14* transgenic plant showed short-narrow flag leaves and thick-green leaves in this study. In *Arabidopsis thaliana*, most miR156-targeted SPL genes have high levels in the shoot apex (Cardon *et al*., 1997; Wu and Poethig, 2006). Moreover, Wang *et al*. (2008) indicated that miR156 quantitatively modulates SPL expression in leaf primordia and SPL mediates nonautonomous effects of existing leaf primordia on the initiation of new leaf primordia at the shoot apical meristem. And plants overexpressing SPL9 showed a strong plastochron with reduced the leaf initiation rate. The research suggested SPLs function in plastochron length and leaf size, but the mechanism including the downstream target gene of SPLs in leaf development is still unknown. Similarly, OsSPL14 was predominantly expressed in the shoot apex (Jiao *et al*., 2010; Miura *et al*., 2010). And OsSPL14 mRNA levels decreased in flag leaves of OsmiR156-overexpression transgenic plants (Xie *et al*., 2006). It suggested the possibility of OsSPL14 involved in development of flag leaf, though there is hardly study on the molecular mechanisms of SPLs in regulating leaf development in rice. Correspondingly, we observed that OE: *OsSPL14* transgenic plants showed greener and the chlorophyll a, chlorophyll b and carotenoid content were higher at seedling stage and at maturity stage. It may chiefly attributable to the up-regulation of genes involved in terpenoid anabolism as description in transcriptome analysis of this study. Nevertheless, there is hardly research on effects of SPLs on leaf color either in rice or in *Arabidopsis thaliana*.

Plant hormones play a vital role in regulating plant growth, development and adaption to environment change (Santner and Estelle, 2009). ABA is a phytohormone that regulates seed dormancy, seed embryo decelopment and stress tolerance improvement (Nambara and Marion-Poll, 2005; Matillaet *al*., 2015). And SLS are hormones involved in suppressing lateral shoot branching, root development, response to environmental condition and secondary growth in vascular plants (Ruyter-Spira, 2013; Al-Babili, 2015). It is notable that ABA and SLs are apocarotenoids shared carotenoid as a biosynthetic precursor (Matusova *et al*., 2005). In accord, our transcriptome analysis revealed that the expression of phytoene synthase gene (*PSY*) and beta-carotene hydroxylase 2 gene (*DSM2*) that the key enzyme genes in carotenoid biosynthesis are up-regulated in OE: *OsSPL14* transgenic plant. Besides, the expression of 9-cis-epoxycarotenoid dioxygenase gene (*NCED*) participated in the rate-limiting step in ABA biosynthesis and genes involved in SLs biosynthesis and SLs signal transduction also changed. Indeed, the carotenoid content of transgenic plant was higher than that of ‘MH86’, and it could provide more precursor for ABA or SLs biosynthesis. According to the common biosynthetic precursor, there is inevitable relation between ABA and SLs. ABA-deficient tomato mutants exhibit greatly reduced ABA and SLs content with the downregulation of SL biosynthetic genes, suggesting a role of ABA in the regulation of SLs biosynthesis (López-Ráez *et al*., 2010). Similarly, SL deficient showed reduced ABA level under osmotic stress (Liu *et al*., 2015). And there is evidence that the biosynthetic pathways of ABA and SLs are functionally connected. Haider *et al*. (2018) indicated that SLs interact with ABA during the drought response in rice. SL and ABA pathways are connected with through the SL biosynthetic enzyme DWARF27 (D27), and overexpression of D27 in rice resulted increased ABA levels. Also, a study indicated that enhanced levels of endogenous ABA cause a decrease in SL production and there was an antagonistic interaction between SLs and ABA biosynthesis in barley. Barley plants expressing an RNAi for *ABA 8’-hydroxylase* involved in ABA degradation had higher ABA content but lower SLs content, and led to increased tiller numbers (Wang *et al*. 2018), but it was likely to occur in the extreme situation. In our study, OE: *OsSPL14* transgenic plants with higher ABA contents still showed decreased tiller numbers. And previous study indicated that OsSPL14 may mediate the SL-regulated tiller development (Song *et al*., 2017). So we conjecture higher ABA could accompany with higher SLs level and they combined effect on plant growth and development.

In addition, gibberellin (GA) is another important growth-promoting hormone. GA is one of tetracyclic diterpenoid plant hormones that participate in diverse growth and developmental processes including leaf differentiation, stem elongation, trichome development, seed germination and induction of flowering (Olszewski N *et al*., 2002; Fleet and Sun, 2005). One main function of GA is regulating organs elongation by affecting cell division and elongation. The gibberellin-deficient dwarf1 (gdd1) rice mutant showed a phenotype of greatly reduced length of root, stems, spikes, and seeds, which can be rescued by exogenous GA_3_ treatment. Further elucidated that GDD1 with transcription regulation activity affects GA accumulation through regulating ent-kaurene oxidase (KO2) involved in GA biosynthesis (Li *et al*., 2011). In Arabidopsis, SPL8, a member of SPL family genes, was proved to be a tissue-dependent regulatory role in response to GA in plant development. And change in SPL8 expression affected transcription of genes involved in GA biosynthesis and signaling (Zhang *et al*., 2006). In this study, we found that the expression of 1-deoxy-D-xylulose-5-phosphate synthase 2 gene (*DXS*) involved in terpenoid anabolism is up-regulated obviously in OE: *OsSPL14* transgenic plant, and the GA_3_ content increased significantly. Previous research showed that rice plant with high OsSPL14 protein level appeared large panicles and increased plant height (Jiao *et al*., 2010; Zhang *et al*., 2017), which may attribute to high GA_3_ level. Moreover, the other main function of GA is promoting flowering. The study indicated that SPL15 was a key component in the integration of age- and GA-derived flowering pathways at the Arabidopsis Shoot Meristem (Hyun *et al*., 2016). Hence, we conjecture that there was a definite relation between early flowering in OE: *OsSPL14* transgenic plant and high GA_3_ level. Based on these, we infer that overexpression of OsSPL14 has influence on several plant hormones biosynthesis so as to regulate plant development. However, the mechanisms are needed to be further elucidated.

In vascular plants, the major component of the secondary cell wall is lignocellulose, which is mainly consisted of lignin, cellulose, hemicellulose, cell wall proteins and so on (Kummar*et al*., 2016). Lignin is a phenylpropanoid monomer polymer that maintains structural integrity of the cell wall and increases mechanical strength of plant. The pathways of lignin monomer biosynthesis start with deamination of phenylalanine by phenylalanine ammonia-lyase (PAL). Then the product cinnamic acid was transferred to coumaric acid by cinnamic acid 4-hydroxylase (C4H), and further acted by 4-coumarate--CoA ligase (4CL) produces *p*-coumaroyl-CoA (Boerjan *et al*., 2003; Zhong and Ye, 2015; Kummar *et al*., 2016). In this study, transcriptome analysis indicated that the expression of PAL, C4H and 4CL were up-regulated in OE: *OsSPL14* transgenic plant. Coincidentally, the lignin content of culm increased dramatically in OE: *OsSPL14* transgenic plant. Additionally, cellulose is a polysaccharide and cellulose microfibrils form the main load-bearing network (Somerville, 2006). Our study suggested that the expression of some genes participated in sugar metabolism changed and the cellulose content of culm increased obviously in OE: *OsSPL14* transgenic plant. Guo *et al*. (2003) indicated that high cellulose and lignin contents within a certain stem enhanced the lodging resistance. Thus, it is clear that higher content of lignin and cellulose would make contributions to enhancement of mechanical strength in *OsSPL14*-overexpression plant. Additionally, quantity of silicon had a close relationship with mechanical strength of culm. Silicon, mainly existed in epidermal layers of stems, leaf sheaths and vascular bundles, can increase strength and hardness of the plant cell walls (Luo *et al*., 2007). The study indicated that increasing silicon content of plants could improve the mechanical strength of the basal internode and eventually increase plant resistance to lodging (Jiang *et al*., 2012). Besides, potassium is another factor impacted on mechanical strength of culm. Yang *et al*. (2004) indicated that thickness of basal stem and anti-fracture correlated with silicon and potassium content. Our result showed that silicon and potassium content increased in OE: *OsSPL14* transgenic plant. It further illustrated that mechanical strength of *OsSPL14*-overexpression plant increased so as to elevate lodging resistance. Together, OsSPL14 is a pleiotropic regulator that participates in several metabolic pathways and plays an important role in growth and development of plants (Fig. 7). Certainly, the regulatory network of OsSPL14 is worth further studying in rice.

**Fig.7.**
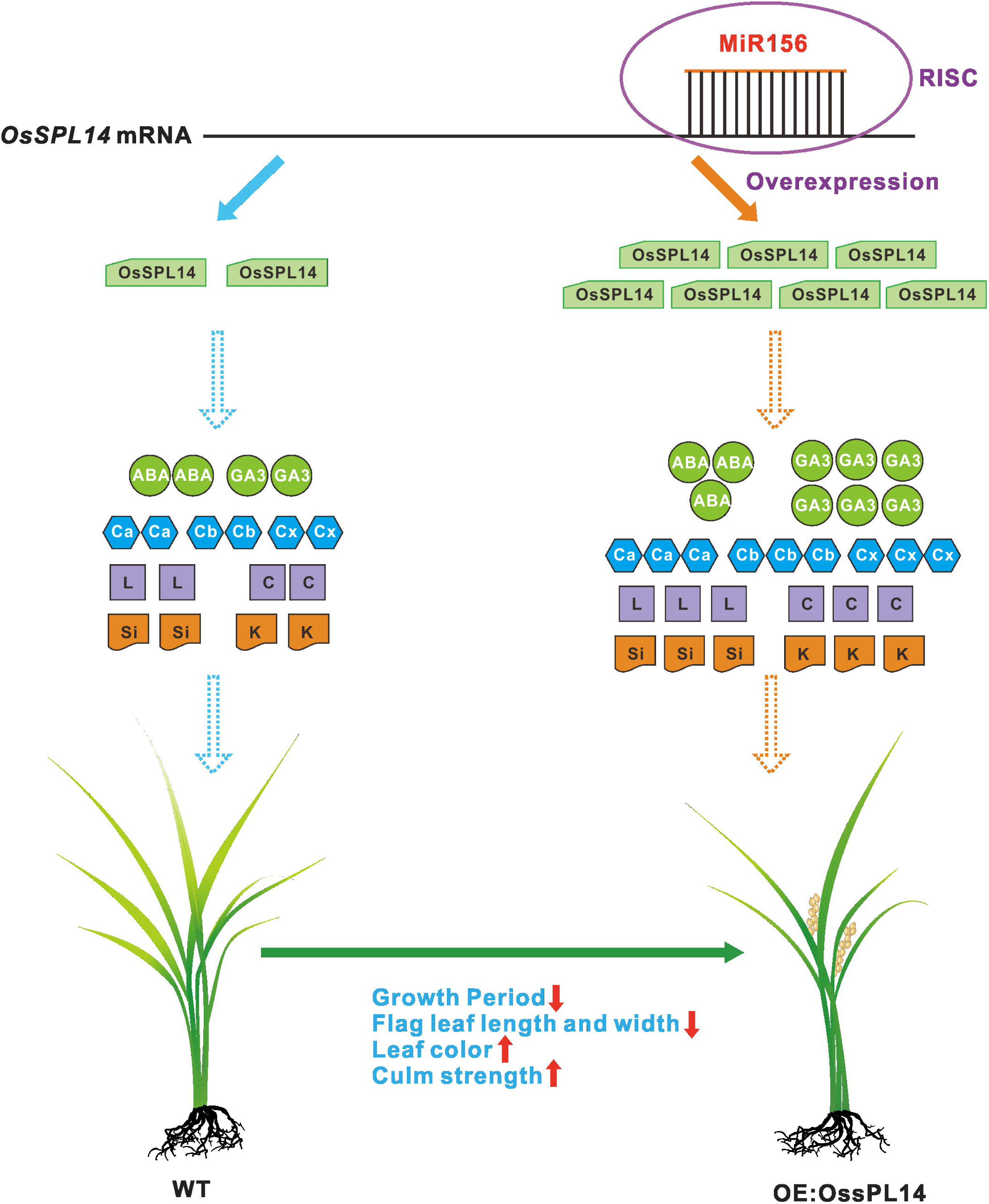
A model for the function of OsSPL14 in regulating plant growth and development. Arrows with dotted line mean there are unclear molecular mechanisms. RISC: RNA-induced silencing complex; L: Lignin; C: Cellulose; Si: Silicon; K: Potassium.

## Supplementary data

**Fig. S1.** The schematic diagram of *OsSPL14* expression vector.

**Fig. S2.** Detection of transgenic plants.

**Table S1.** Primers used in this study.

**Table S2.** KEGG enrichment analysis of DEGs.

## Acknowledgements

The plasmid pCAMBIA1300-*OsSPL14* was provided by Professor Jiayang Li, the Institute of Genetics and Developmental Biology, Chinese Academy of Science, China. And we thank Prof. Jiayang Li for reading the manuscript. This work was supported by the Key Program of National Transgenic Research of China (Grant no. 2016ZX08001-004) and The Special Foundation of Non-Profit Research Institutes of Fujian Province (Grant no. 2016R1020-8).

## Abbreviations

ABA: abscisic acid
DEGs: differentially expressed genes
FC: fold Change
FDR: false discovery rate
GA_3_: gibberellin acid
GO: gene ontology
JA: jasmonate acid
KEGG: kyotoencyclopedia of genes and genomes
qRT-PCR: quantitative real-time PCR

